# Molecular architecture of the human caveolin-1 complex

**DOI:** 10.1101/2022.02.17.480763

**Authors:** Jason C. Porta, Bing Han, Alican Gulsevin, Jeongmin Chung, Yelena Peskova, Sarah Connolly, Hassane S. Mchaourab, Jens Meiler, Erkan Karakas, Anne K. Kenworthy, Melanie D. Ohi

## Abstract

Membrane sculpting proteins shape the morphology of cell membranes and facilitate remodeling in response to physiological and environmental cues. Complexes of the monotopic membrane protein caveolin function as essential curvature-generating components of caveolae, flask-shaped invaginations that sense and respond to plasma membrane tension. However, the structural basis for caveolin’s membrane remodeling activity is currently unknown. Here, we show, using cryo-electron microscopy, that the human caveolin-1 complex is composed of 11 protomers organized into a tightly packed disc with a flat membrane-embedded surface. The structural insights suggest a new mechanism for how membrane sculpting proteins interact with membranes and reveal how key regions of caveolin-1, including its scaffolding, oligomerization, and intramembrane domains, contribute to its function.

**One-Sentence Summary:** Cryo-electron microscopy reveals that Caveolin-1 oligomerizes into a tightly packed disc with a flat membrane-binding surface.

## Main Text

Biological membranes assume a wide range of morphologies and are actively remodeled to support the normal physiological functions of cells as well as cell growth and differentiation (*1*, *2*). An important mechanism for shaping and remodeling membranes is through the activity of curvature-inducing proteins (*2*–*4*). Caveolins, a family of monotopic membrane proteins first identified over 30 years ago, fall into this class of proteins based on their ability to bend membranes to form 50-100 nm flask shaped invaginations of the plasma membrane known as caveolae (*5*, *6*). Comprising up to 30-50% of the surface area of some cell types (*7*), caveolae serve as mechanosensors and mechanoprotectors that sense and respond to changes in membrane tension at the cell surface by reversibly flattening (*8*). The absence of caveolin-1 (Cav1) and thus caveolae has profound consequences for mice, including decreased lifespan, lipid metabolism disorders, vascular abnormalities, dilated cardiomyopathy, and pulmonary hypertension (*9*). In humans, Cav1 has been linked to cardiovascular disease, cancer, lipodystrophy, and kidney disease (*10*–*15*). How Cav1 facilitates caveolae biogenesis is not fully understood, but is thought to require the insertion of a highly hydrophobic region of the protein, termed the intramembrane domain (IMD), into the cytoplasmic face of the plasma membrane to induce membrane curvature via a wedging mechanism (*3*, *4*, *16*, *17*). To induce curvature, Cav1 must also assemble correctly into oligomeric complexes 8S in size that undergo additional higher order interactions with each other and the cavin proteins to form mature caveolae that are polygonal in shape (*18*–*21*). 8S complex formation occurs via a cooperative process mediated by its oligomerization domain (OD) and aided by its scaffolding domain (SD) and signature motif (SM) (*21*–*23*). However, the structural basis for Cav1’s membrane bending activity remains unknown, as only low-resolution structures of caveolin complexes currently exist (*24*–*26*).

Although caveolin proteins are found exclusively in metazoans (*18*), when expressed in *Escherichia coli*, Cav1 forms 8S-like complexes (*26*) and induces the formation of heterologous caveolae (*h*-caveolae) (*27*, *28*). Thus, *E. coli*-expressed Cav1 forms oligomers and sculpts membranes, two of its essential functions in mammalian cells. Using single-particle cryo-EM analysis, we determined the structure of *E. coli* expressed and purified complexes (*26*) (Fig. 1, Figs. S1-3, and Methods). 2D class averages of *en face* views of the complex showed 11 spiraled α -helices, and an *ab initio* 3D reconstruction with no enforced symmetry resulted in a low-resolution map with structural features indicating potential 11-fold symmetry (Fig. S3C, D). Applying C11 symmetry during 3D refinement steps led to a 3.5 Å resolution structure (Fig. 1 and Fig. S4). This agrees with previous reports suggesting 7-14 copies of Cav1 per 8S complex (*21*, *22*, *26*). The 11 Cav1 protomers organize into a disc-shaped complex with a diameter of ~140 Å and a height of ~34 Å (Fig. 1A-F). To ensure we were not missing other oligomeric forms of the complex by enforcing 11-fold symmetry, we performed 3D refinements using symmetries spanning C8-C14. None of these resulted in maps with apparent secondary structural features or improved resolutions (Fig. S5). Additionally, 3D classification did not yield models of any other oligomeric forms.

**Figure 1.**
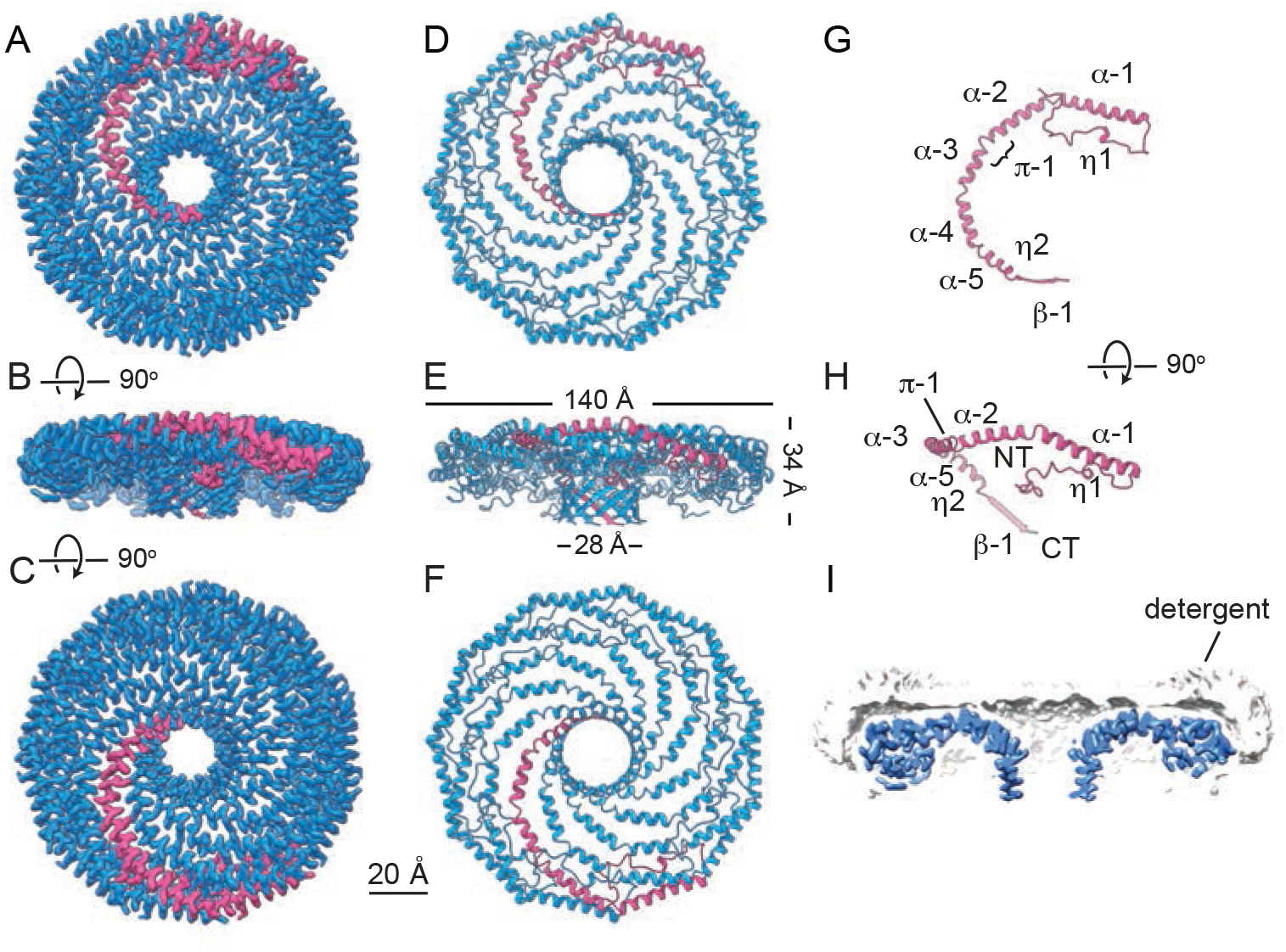
Cryo-EM structure of the 8S Cav1 complex. **(A-C)** 90° rotated views of the cryo-EM density of the 8S Cav1 complex at 3.5 Å with 11-fold symmetry. The complex is a disc-shaped structure composed of tightly packed a-helices and a cylindrical β-barrel. Magenta, Cav1 protomer. **(D-F)**. Secondary structure model of the refined 8S Cav1 complex. Same views as shown in A-C. Dimensions of complex are labeled in E. **(G-H)** Secondary structure of Cav1 protomer with secondary structural features and position of the N- and C-termini labeled. **(I)** Central slice of the density map (blue) with detergent micelle (light grey). Scale bar for all panels, 20Å.

The structure reveals Cav1 oligomerizes into a disc that contains an outer “rim” (~23 Å wide), a central β-barrel “hub” (~28 Å wide), and eleven curved a-helical “spokes”. Each Cav1 protomer is mostly α-helical (α1-5), consistent with previous reports (*29*) (Fig. 1D-H). Viewed from the side, helices α2-α5 of the protomer align along the flat surface of the complex, and helix 1 forms a ~20° angle to this surface (Fig. 1H). Cav1 also contains two non-helical regions located at both termini (Fig. 1D-H). The N-terminus, which makes up part of the “rim” region, consists of a loop with a short 3_10_ helix (η1) (Fig 1G, H). This region makes a 180° turn at the outer edge of the complex, intercalating with the spokes and N-terminal regions of neighboring protomers (Fig. 1A-H). Each protomer terminates in a C-terminal β-strand angled by ~40° relative to the plane of the flat surface of the disc. These β-strands interact to form an 11-stranded parallel β-barrel “hub” (Fig. 1A-H). Based on the position of the detergent micelle in the density map (Fig. 1I) and our previous negative stain analysis of *h*-caveolae (*26*), the flat face of the complex corresponds to the membrane-facing surface, whereas the β-barrel faces the cytoplasm. Strikingly, the central cavity of the barrel extends entirely through the complex and is open to solvent at the cytoplasmic-facing side (Fig. 1A-F and 1I). The detergent micelle covers ~60% of the surface area of the 8S complex, including the membrane facing surface, and extends ~14Å around the sides of the outer rim of the disc (Fig. 1I).

From the density map, we built an atomic model for most of Cav1 (residues 49-177) (Fig. 2A-D). However, residues 1-48, which fall within a predicted disordered region (*28*, *29*) and are dispensable for caveolae biogenesis (*18*), were not detected in the map (Fig. 2A-D and Fig. S6). To correlate the extensive biochemical and functional data in the literature with our structural analysis, we mapped key regions and residues of Cav1 onto the structure (Fig. 2A-G, Fig. S7,and Supplementary Video 1). The OD (residues 61-101) (*22*) is located at the outer rim of the disc and contributes to extensive subunit interfaces. The SM (residues 68-75) (*30*, *31*) and SD (residues 82-101) (*32*) both fall within the OD. Lying within the loop region, the SM forms tight contacts with two neighboring protomers, whereas the SD comprises most of helix α1 and encircles the periphery of the complex. The residues separating the SM and SD make two 90° turns, bringing these two motifs into proximity (Fig. 2D). The IMD (residues 102-134) (*23*, *29*) begins on the C-terminal end of the helix α1 immediately adjacent to the SD and continues across helices α2, α1, and part of α3 (Fig. 2A-D). Known phosphorylation and ubiquitination (*33*, *34*) sites are accessible on the cytoplasmic face of the complex, whereas Cav1’s palmitoylation sites (Cys133, Cys143, and Cys156) (*35*) are located on the membrane-facing surface (Fig. S8).

**Figure 2.**
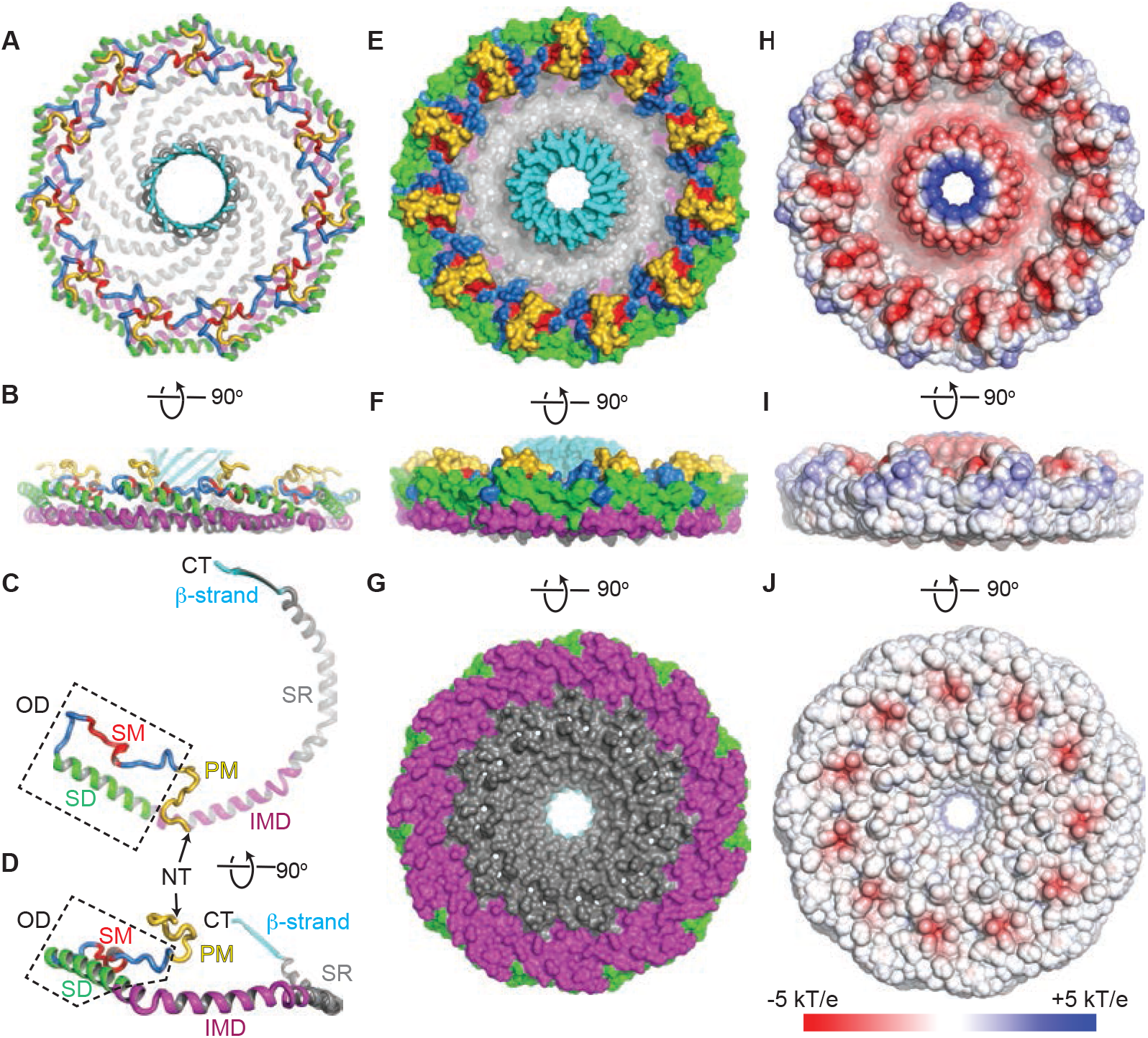
Structure of Cav1. (**A**)3.5 Å resolution structure of the 8S Cav1 complex as viewed from the cytoplasmic face. (**B**) Structure rotated 90°. (**C-D**) Structure of Cav1 rotated 90°. The positions of previously defined regions are labeled: SM, signature motif (red); SD, scaffolding domain (green); and IMD, intermembrane domain (purple). The OD, which contains the SM and SD, is indicated by the dashed box. New structurally defined motifs include: PM, pin motif (yellow), SR, spoke region (grey); and β-strand (cyan). N-terminus (NT) and C-terminus (CT) are marked with arrows. (**E-G**) Space filling model of the 8S Cav1 complex rotated 90°. Color scheme identical as in A-D. (**H-J**) Space filling model of the 8S Cav1 complex rotated 90° showing the charge of the amino acids. Red, negative; grey, neutral; blue, positive.

In addition to these previously characterized functional regions, we identified three new motifs in Cav1 that are critical for 8S complex stability. First, residues 49-60 form a loop extending over the SM of the neighboring protomer (Figs. 2A-D and 3). This previously unnamed region is required for caveolae assembly (*18*) and localization in migrating fibroblasts (*36*). The structure shows that these residues appear to “lock” the interaction between protomers, leading us to designate this region as the “pin motif” (PM) (Fig. 2A-G and 3; Fig. S7). Second, residues 135-169, a region required for caveolae formation (*18*), oligomerization, and exit of the protein from the Golgi complex (*20*), generate a spoke region (SR) organized parallel to the membrane plane (Fig. 2A-D; Fig. S7). The structure shows these residues form the planar hydrophobic surface on the membrane-facing side of the complex while also creating a highly negatively charged surface on the cytoplasm-facing side of the complex (Fig. 2G, J). Helix α5 (residues 143-155) distorts this flat surface by bending towards the cytoplasm facing side of the complex where it connects to a β-strand (residues 170-176) (Figs. 1G, H and 2C, D). β-strands from adjacent protomers assemble into the third new motif, an eleven-stranded parallel β-barrel with a hydrophilic exterior and hydrophobic interior (Fig. 2A-G). To our knowledge, this represents the largest reported example of a parallel β-barrel to date; the most similar β-barrel reported in the PDB (2AO9) was observed in a soluble bacteriophage protein of unknown function and contains nine parallel β-strands (Fig. S9A). The barrel is capped by Lys176, introducing a highly positively charged layer separating the hydrophobic interior of the barrel from the cytoplasm. The narrowest accessible region of the interior of the β-barrel has a diameter of 15 Å (Fig. S9B), making the channel large enough to accommodate small molecules such as lipids. However, no density was detected inside the β-barrel, perhaps because of the imposed C11 symmetry.

**Figure 3.**
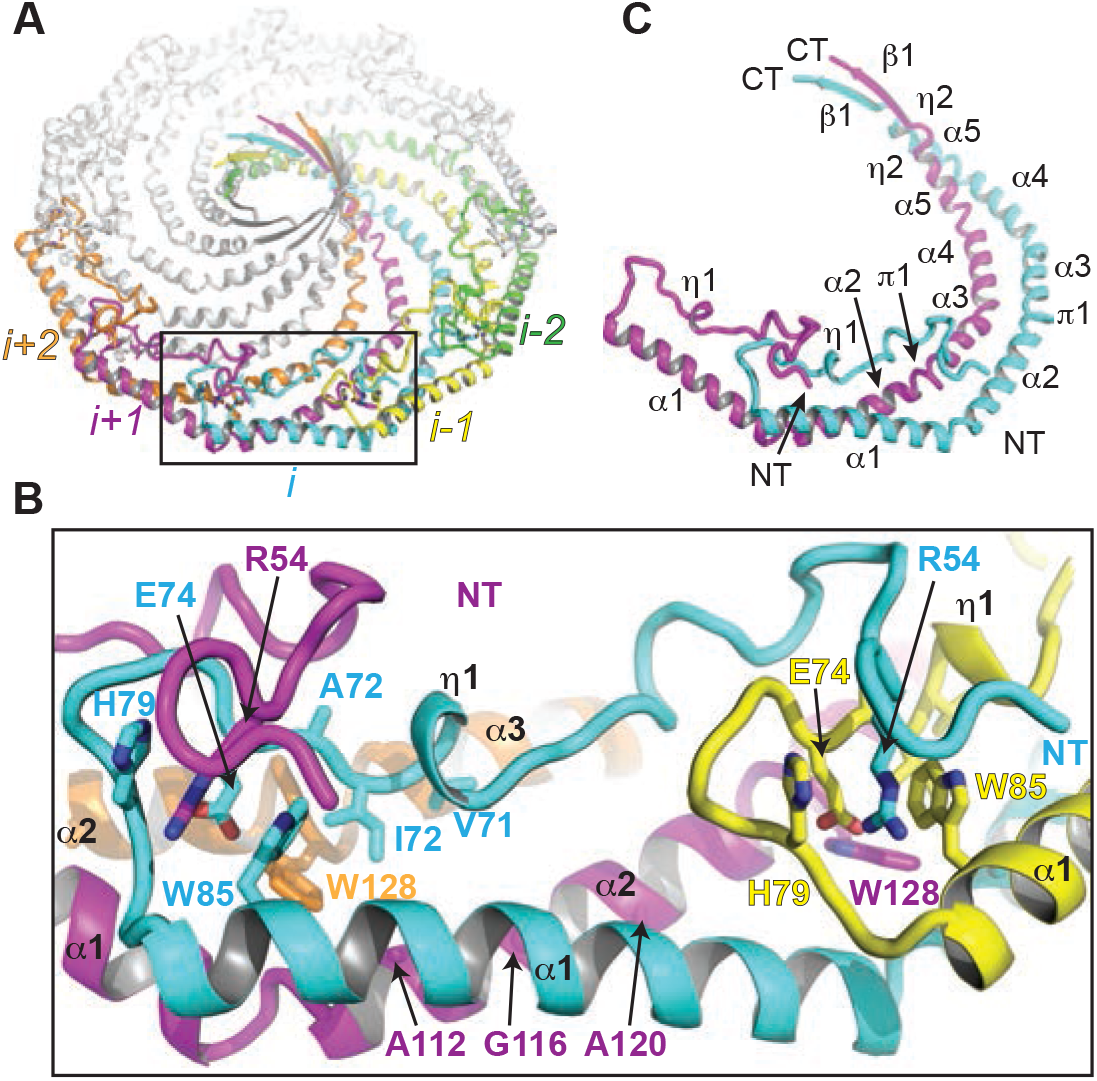
Cav1 8S complex is stabilized by extensive interactions along the length of the protomers. (**A**) Overall structure of the 8S complex highlighting 5 distinctly colored protomers labeled i-2 to i+2. (**B**) Zoomed view of the rim region (box in panel A) highlighting key interacting residues. (**C**) Packing of two protomers. Secondary structure elements are labeled.

The structure also unveils atomic details of how protomers oligomerize to form 8S complexes, a process essential for caveolae biogenesis. Classically, the OD has been proposed to function as the main region driving protomer-protomer interactions that lead to 8S complex formation (*9*). The structure reveals that instead extensive networks of interactions occur along the entirety of each protomer, including the rim, spoke, and hub regions (Fig. 3 and Fig. S9C-F). Starting at the “rim”, the N-terminus of one protomer (*i*) engages four neighboring protomers (*i*-2, *i*-1, *i*+1, and *i*+2) (Fig. 3A, B). The ODs from neighboring protomers are arranged adjacent to each other, but only slightly overlap (Fig. 3B). Instead, the OD of protomer *i* stacks onto the cytoplasmic side of the IMDs of the neighboring protomers *i*+1 and *i*+2 (Fig. 3A, B). Within OD_*i*_, SM_*i*_, is sandwiched between α1_*i*+1_, α2_*i*+1_, α2_*i*+2_, α3_*i*,+2_, π1_*i*+2_ and PM_*i*+1_. Arg54_*i*+1_ of PM_*i*+1_ intercalates into a pocket formed by His79_*i*_, and Trp85_*i*_, pinning OD_*i*_, in place (Fig. 3A,B). To test the importance of this newly described molecular “pin”, we mutated Arg54, a residue predicted to play an important role in facilitating protomer-protomer interactions, to alanine. This mutation severely distrupts the 8S complexes, confirming that the PM interaction with the SM of the neighboring protomer is important for the 8S complex formation (Fig. S10). The highly hydrophobic IMD helices (α2, π1, and part of α3) lay underneath the OD of the neighboring protomers, contributing to oligomerization (Fig. 3A, C). α2_*i*+2_ forms a helical bundle that crosses α1_*i*+1_ facilitated by the small side chains of residues Ala112_*i*+1_, Gly116_*i*+1_, and Ala120_*i*+1_. The side chain of Trp128_*i*+2_ extends from π1_*i*+2_ into the cavity formed by OD_*i*_, and the loop between α1_*i*+1_ and α2_*i*+1_, suggesting it plays a critical role in oligomerization (Fig. 3A, C). The remaining helices forming the SR (C-terminal portion of α3, α4, and α5) make pairwise contacts across protomers (Fig. 3C, Fig. S9C-F). These interactions involve residues with larger side chains, increasing the separation of the helices compared to those of the α1_*i*_, -α2_*i*+1_ crossing, which ultimately converge to form the C-terminal β-barrel, the final major region of protomer-protomer interactions. The symmetry dictates a two-residue offset between neighboring protomers, creating a β-barrel (11, 22). Interestingly, several disease-associated mutations in Cav1 localize on the connecting regions between the IMD and SR, or the SR and β-barrel (Fig. S8), suggesting that this region of the complex is sensitive to mutations that destabilize the complex (*14*, *15*).

The structure of the human Cav1 8S complex provides a molecular framework for understanding the overall organization of Cav1 oligomers, the exact roles the OD, SD, SM, and IMD play in complex formation, the importance of previously unrecognized regions of the protein, and the impact of disease-associated mutations such as P132L (*12*) (Fig. S8H, I) on the structure. Importantly, it also suggests new models for how Cav1 packs in membranes and controls caveolae architecture (Fig. S11). In contrast to the prevailing model suggesting the IMD forms a hairpin-like structure that inserts into the membrane creating a wedge that bends membrane (*3*, *4*, *16*, *17*), the structure of the 8S Cav1 complex reveals that the IMD contributes to the formation of a flat membrane-facing surface while simultaneously stabilizing contacts between protomers. The outside of the outer rim of the complex is also primarily hydrophobic. Together, these features suggest a model in which the membrane-associated side of the complex embeds deeply within the cytoplasmic leaflet, interacting with the terminal carbons of the lipids of the opposing leaflet rather than sitting at the interface between the headgroups and acyl chains as typically would be expected for amphipathic helices (Fig. 4B). In this model, the 8S complex, by displacing lipids from the cytoplasmic leaflet, could create an ordered membrane nanodomain composed of protein on one membrane leaflet and lipids on the other. Interestingly, a single 8S complex fits snugly on each face of a dodecahedron of the characteristic size of *h*-caveolae (30 nm average diameter) (*28*) (Fig. S11a), whereas up to three can be accommodated on each surface of a mammalian caveolae (61 nm average diameter) assuming dodecahedral symmetry (*25*) (Fig. S11B-C). Thus, in caveolae, caveolin complexes may function primarily by stabilizing flat membrane surfaces of polyhedral structures rather than imposing continuous membrane curvature, defining a new mechanism for how integral membrane proteins sculpt cell membranes to form functional domains.

**Figure 4.**
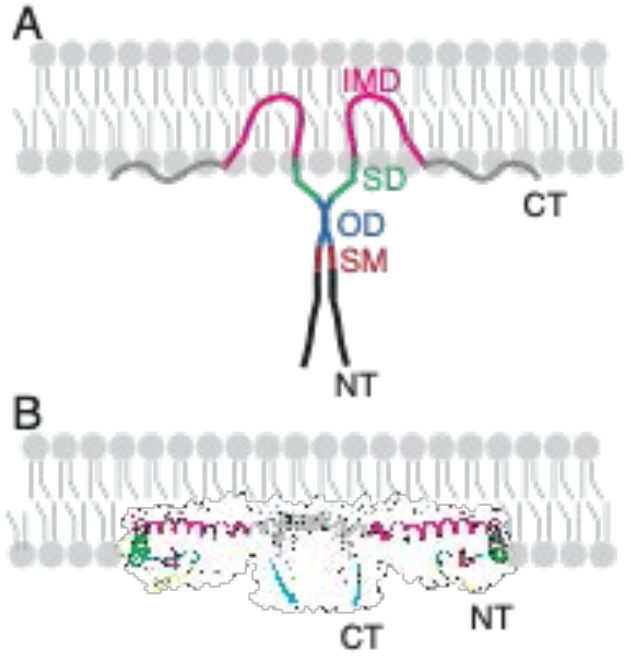
Proposed model for Cav1 8S complex association with membranes. (**A**) Classical model of Cav1 oligomer organization and membrane interaction.(**B**) Structure-based model. See text for details. Coloring scheme matches Figure 2A-D.

## Acknowledgments

We thank Dr. K. Jebrell Glover for providing cDNA constructs and Drs. Jochen Zimmer, Charles Sanders, James Casanova, and Ludger Johannes for feedback on the manuscript. The University of Michigan Cryo-EM Facility (U-M Cryo-EM) has received generous support from the U-M Life Sciences Institute and the U-M Biosciences Initiative.

## Author contributions

Conceptualization: AKK, MDO, EK, HSM, JM

Methodology: JCP, JC, BH, YP

Investigation: JCP, BH, YP, JC, AG, SC

Data curation: JCP, JC

Formal analysis: JCP, AG, JC, EK

Writing-original draft: AKK, MDO, EK

Writing-review and editing: JCP, BH, AG, JC, YP, SC, HSM, JM, EK, AKK, MDO

Visualization: JCP, BH, AG, SC, HSM, EK, MDO

Supervision: AKK, MDO, EK, JM

Project administration: AKK, MDO

Funding acquisition: AKK, MDO, JM.

## Data and materials availability

The cryo-EM volume and the structure coordinates have been deposited in the Electron Microscopy Data Bank and the Protein Data Bank under accession codes EMD-XXXX and XXXX.

## Supplementary Materials

Materials and Methods

Figs. S1 to S11

Tables S1

References (*37*–*62*)

