## Supplementary Material for "Molecular architecture of the human caveolin-1 complex"

**This PDF file includes:**

Materials and Methods

Figs. S1 to S11

Tables S1

References (*37*–*62*)

### Materials and Methods

**Construct preparation.** Cav1-MycHis plasmid was constructed by site-directed mutagenesis using QuikChange Lightning Kit (Agilent, #210518). The template was Cysteine-less Cav1-MycHis (a gift from K. Jebrell Glover, Lehigh University). Primers were used to introduce native cysteines in the positions 133 (primer: GGCGGTTGTTCCGTGCATCAAATCTTTCCTGATCG), 143 (primer: CCTGATCGAAATCCAGTGCATCTCTCGTGTTTACTC) and 156 (primer: CGTTCACACCGTTTGCGACCCGCTGTTCG). Gene sequences were confirmed by sequencing (Genewiz, NJ).

**Expression and purification of the 8S Cav1 complex.** Protein expression and purification were conducted as described before (*26*) with minor modifications. Cav1 was expressed in *E. coli* BL21 using the auto-induction expression system (*37*). The MDG starter of monoclonal bacteria cultured at 37°C and 250 rpm for 20 hours was transferred into the auto-inducing ZYM-5052 media and incubated at 25°C and 300 rpm for 24 hours. The harvested cells were first washed with 0.9% NaCl and then resuspended in a buffer composed of 200 mM NaCl and 20 mM Tris-HCl (pH 8.0), 1mM PMSF, and 1mM DTT. The cell suspension was homogenized using a French press pressure homogenizer and centrifuged at 9,000 rpm for 15 min. at 4°C to remove large cell debris. The supernatant was then centrifuged at 40,000 rpm (Ti-45 rotor) to pellet the membrane fractions. Membrane pellets were homogenized in a buffer composed of 200 mM NaCl, 20 mM Tris-HCl (pH 8.0), and 1 mM DTT using a Dounce tissue grinder. Solubilization of the membrane was performed by adding n-Dodecyl β-D-maltoside (C12M, Anatrace, Ohio) to a final concentration of 2%, and incubating with gentle agitation for 2 hours at 4°C. After centrifugation at 42,000 rpm (Ti-50.2 rotor) for 35 min to remove insoluble material, the supernatant was used for nickel sepharose–based affinity purification. Specifically, the supernatant was incubated with nickel-charged Sepharose (Chelating Sepharose Fast Flow, GE

Healthcare, Illinois) that was pre-equilibrated with buffer of 200 mM NaCl, 20 mM Tris-HCl (pH 8.0), 0.05% C12M, 10mM imidazole and 1 mM DTT for 2 hours at 4°C. After incubation, the resin–supernatant mixture was poured into a column and was washed with 30mM imidazole in the above-described buffer. The protein was then eluted using the buffer supplemented with 300 mM imidazole. The elutions with caveolin proteins were concentrated and further separated by size exclusion chromatography by using Superpose 6 Increase 10/300 GL column (GE Healthcare, Illinois) equilibrated with a buffer composed of 200 mM NaCl, 20 mM Tris-HCl (pH 8.0), 1 mM DTT, and 0.05% C12M. Similar to our previous report (*26*), two major peaks were observed in the SEC profile, both of which were enriched with Cav1 proteins (Fig. S1). The P1 fractions containing the fully oligomerized 8S complexes were diluted from ~1.1 mg/ml to a final concentration of ~0.11 mg/ml for cryo-EM experiments.

**Electrophoresis and Western blotting.** SDS-PAGE Electrophoresis and Western blotting were performed as described previously (*14*). Mouse anti-Cav1 polyclonal antibody (Catalog number: 610406, BD Biosciences, San Jose, CA) and rabbit anti-His-Tag (Catalog number: A1138-50, BioVision, California) was used with 1:2,000 times dilution. Secondary antibodies and blocking buffer were obtained from LI-COR and imaging was performed using the LI-COR Odyssey system (LI-COR Biosciences, Nebraska).

**Negative Stain imaging and data processing**. Negative stain EM was done using established methods (*38*). In brief, 200-mesh copper grids covered with carbon-coated collodion film (EMS, Hatfield, PA, USA) were glow discharged for 30 s at 10 mA in a PELCO easiGlow™ glow discharge unit (Fresno, CA, USA). Aliquots (3.5 μl) of purified CAV1-myc-his (50 μg/ml) or Cav1-R54A-myc-his (30 μg/ml) were adsorbed to the grids and incubated for 1 minute at room temperature. Samples were then washed with 2 drops of water and stained with 2 successive drops of 0.7% (w/v) uranyl formate (EMS, Hatfield, PA, USA) followed by blotting until dry. Samples were visualized on a Morgagni transmission electron microscope equipped with a field emission gun operating at an accelerating voltage of 100 keV (Thermo Fisher Scientific Inc., Waltham, MA, USA) at a nominal magnification of 22,000x (2.1 Å per pixel). The Cav1-R54A-myc-his negative stain dataset was collected using a Tecnai Spirit T12 transmission electron microscope operated at 120keV (Thermo Fisher Scientific, Waltham, MA, USA) at a nominal magnification of ×26,000 (2.34 Å per pixel). Sample data were collected using Leginon software on a 4 k × 4 k Rio complementary metal-oxide semiconductor camera (Gatan, Pleasanton, CA) at −1.5-µm defocus value. Images were manually curated. All data processing was performed in Relion 3.1.0. About 1,000 particles were picked manually and 2D classified. Clear resulting classes were selected as references for particle selection on all images. Particles were extracted with a 128 pixel box size (30nm by 30nm) and then underwent 2D classification. The Cav1-R54A-myc-his dataset consisted of 54,874 particles.

**Cryo-EM sample preparation.** For cryo-EM, 4 µl of the protein sample was applied to Quantifoil 400-mesh Au grids (Electron Microscopy Sciences) that were glow discharged for 30 s at 5 mA. The sample was then vitrified by plunge freezing in a liquid ethane slurry using a Vitrobot Mark IV robotic plunger (Thermo Fisher). The chamber was set to 100% humidity, and sample grids were blotted for 5 s with a blot force of 10.

**Cryo-EM data collection.** All images were collected on a Glacios transmission electron microscope (Thermo Fisher) operated at 200 keV and equipped with a K2 Summit direct electron detector (Gatan) at a nominal pixel size of 0.98 Å. The total exposure time was 6 s, and frames were recorded every 0.2 s, giving an accumulated dose of 55.5 $e^{-}$ $Å^{-2}$ using a defocus range of -1.5 µm to -2.2 µm. Images were acquired using Leginon software (*39*).

**Image processing.** All image processing, classification, and map refinements were done in cryoSPARC (*40*). After image acquisition, 984 movie frames were dose-weighted and locally corrected for beam-induced drift using the patch motion correction. The contrast transfer function (CTF) parameters were estimated locally over each micrograph using the patch CTF estimation procedure. We first performed non-template-based particle picking on 100 images with the Blob Picker. The resulting 47,892 particles were used as input into the Topaz Train routine (*41*) to generate a neural network model of the particle data. Following the training procedure, the particle model was then input into the Topaz Extract routine over all micrographs, which identified 176,936 particle locations. Particles were extracted into 400 $px^2$ boxes (392 Å x 392 Å). The number of the particles dropped to 140,242 after discarding particles at the micrograph edges. Following two successive rounds of 2D classification, 95,261 particles were selected and used for *ab initio* reconstruction, searching for two classes. One of the classes consisting of 60,615 particles produced a low-resolution map with features agreeing with the previous negative stain EM studies (*25, 26*). The *ab initio* reconstruction of Cav1 along with several 2D class averages displayed a C11 symmetrical structure of the 8S complex (Fig. S3). For this reason, the *ab initio* reconstruction and its associated particles were then submitted to the non-uniform refinement procedure, followed by local refinement using a mask encapsulating the protein only and imposing C11 symmetry (*40*). The masked FSC resolutions from both non-uniform and local refinement were 3.8 Å and 3.5 Å, respectively. The final refined B-factors from the non-uniform and local refinements were 174.5 $Å^2$ and 154.6 $Å^2$, respectively. Gold standard FSC calculations using the Fourier Shell Correlation (FSC) Server (EMBL-EBI) of the local refinement half-maps indicated a global resolution of 3.5 Å at a 0.143 cutoff (Table S1).

**Model building, refinement, and validation.** The density map of the 8S Caveolin complex was of sufficient quality for building a *de novo* model of protein. The regions with well-resolved secondary structures (residues 80-175) for one of the protomers were built using iterative cycles of DeepTracer (*42*), RosettaRelax (*43, 44*), and RosettaES (*45*). Specifically, we first segmented the map manually to obtain the putative monomeric region that corresponds to $Cav1_{60-178}$. The initial prediction of the amino acid coordinates was performed with DeepTracer (*42*) using the human Cav1 sequence range 70 to 178. Next, the model generated by DeepTracer was relaxed with RosettaRelax (*43, 44*) using cryo-EM densities as restraints to build the side chains. Two helical regions that had the most distinctive cluster of aromatic amino acids based on the sequence ($Cav1_{89-107}$ and $Cav1_{111-130}$) were built manually as ideal α-helices. The modeled structures were relaxed into the density domains corresponding to these regions with RosettaRelax with cryo-EM restraints. The two helices were combined in a single PDB file and this file was used as the template for the following RosettaES (*45*) calculations. RosettaES was used to build the $Cav1_{80-178}$ model in two steps. First, the input consisting of the two helices was used as the template to build the three missing segments of the structure with predominantly helical secondary structures ($Cav1_{80-89}$, $Cav1_{108-110}$, and $Cav1_{130-163}$). Second, the input from the first RosettaES modeling step was used to build the residues between 163 and 178 that have less well-defined secondary structures using the same parameters. The resulting structure corresponded to $Cav1_{80-175}$.

The remainder of the model (residues 49-79) was built manually using Coot (V0.9.5) (*46*). The model for the protomer was expanded into undecamer using the C11 symmetry, followed by the refinement of the model using Phenix Real Space Refinement (*47*) and Phenix-Rosetta (*48*). We repeated build-refine iterations until a satisfactory model was obtained. The resolution of the model was estimated by Fourier Shell correlation against the map used to construct it within the Phenix Cryo-EM Validation tool. Molprobity scores, Clashscores, and Ramachandran plots were calculated using Molprobity (*49*) (Extended Table 1). Programs used for structure determination and refinement were accessed through SBGrid (*50*). Figures were prepared using Chimera (v1.15) (*51*), ChimeraX (*52*), and PyMol (v2, Schrödinger, LLC).

**Search for structurally homologous proteins.** Four servers were used to investigate similar motifs to that of caveolin-1 using $CAV1_{49-178}$ structure as the input: the PDBe Fold server (*53*), DALI server (*54*), RUPEE server (*55*), and FATCAT server (*56*). In addition to these analyses, membrane protein structures from the OPM database (*57*) were investigated. 508 entries classified as "bitopic membrane proteins" and 1574 entries classified as "monotopic/peripheral" were visually inspected to identify motifs similar to caveolin-1. The only match found by the PDBe Fold server was toxofilin (2Q97.T) (*58*) with a Q value of 0.051. When the sequences were compared with pBLAST no similarity was found even at a threshold level of 1000, suggesting the scaffold similarity is not related to the amino acid sequence. The DALI server search results yielded long α-helices or helix-turn-helix motifs, but visual inspection of these hits showed no other similarity to the unique structural motifs of caveolin-1. The RUPEE server search resulted in 400 structures, but none of these structures had TM-score values above 0.5 that is accepted as a cutoff to consider two structures similar. The FATCAT server search resulted in 4,222 hits, but almost all structures consisted of a single long α-helix with no sequence similarity. The exception was 6VQ6.M (*59*), which showed two helices connected by a break and a coiled C-terminus, but it had no sequence similarity to caveolin-1.

The SCOP database (*60, 61*) was manually inspected through a keyword search for proteins that have barrel motifs. Of the resulting 89 families, the 2AO9 entry that belongs to the "BC1890-like family" showed similarity to the caveolin-1 structure in terms of the presence of a parallel β-barrel formed by multiple protomeric units, and thus was considered for our structural comparisons.

**Extended Data Table 1 | Cryo-EM data collection, refinement and validation statistics**

| | Caveolin-1 8S complex (EMDB-XXXX) (PDB XXX) |
|---|---|
| **Data collection and processing** | |
| Magnification | 45,000 |
| Voltage (kV) | 300 |
| Electron exposure (e–/Å$^2$) | 55.5 |
| Defocus range (μm) | 1.2-3.2 |
| Pixel size (Å) | 0.98 |
| Symmetry imposed | C11 |
| Initial particle images (no.) | 176,396 |
| Final particle images (no.) | 60,615 |
| Map resolution (Å) | 3.5 |
| FSC threshold | 0.143 |
| Map resolution range (Å) | 3.2-3.7 |
| **Refinement** | |
| Initial model used (PDB code) | None |
| Model resolution (Å) | 3.7 |
| FSC threshold | 0.5 |
| Map sharpening *B* factor (Å$^2$) | 154.6 |
| Model composition | |
| Non-hydrogen atoms | 11,638 |
| Protein residues | 1,419 |
| Ligands | None |
| *B* factors (Å$^2$) | |
| Protein | -100 |
| Ligand | None |
| R.m.s. deviations | |
| Bond lengths (Å) | 0.002 |
| Bond angles (°) | 0.432 |
| Validation | |
| MolProbity score | |
| Clashscore | 1.29 |
| Poor rotamers (%) | 0 |
| Ramachandran plot | |
| Favored (%) | 92.13 |
| Allowed (%) | 7.87 |
| Disallowed (%) | 0 |

**Supplementary Figures**

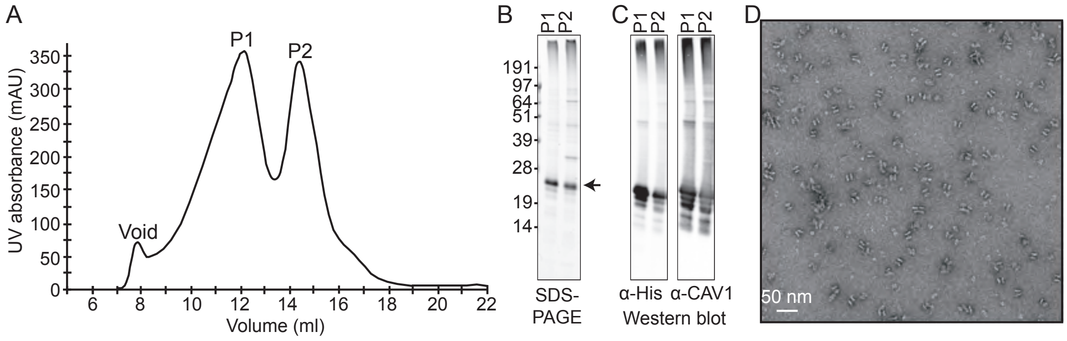

**Fig. S1. Purification of the 8S Cav1 Complex from *E. coli*. (A**) Gel filtration profile of detergent solubilized Cav1-myc-his oligomers. The position of the void and peaks P1 and P2 are labeled. mAU, milli-absorbance unit. **(B)** Coomassie-stained SDS-PAGE gel of P1 and P2. Arrow marks the position of Cav1. **(C)** Western blot analysis shows that each peak is composed of Cav1. **D**. Representative negative stain image of particles from P1 shown in a. Scale bar, 50 nm.

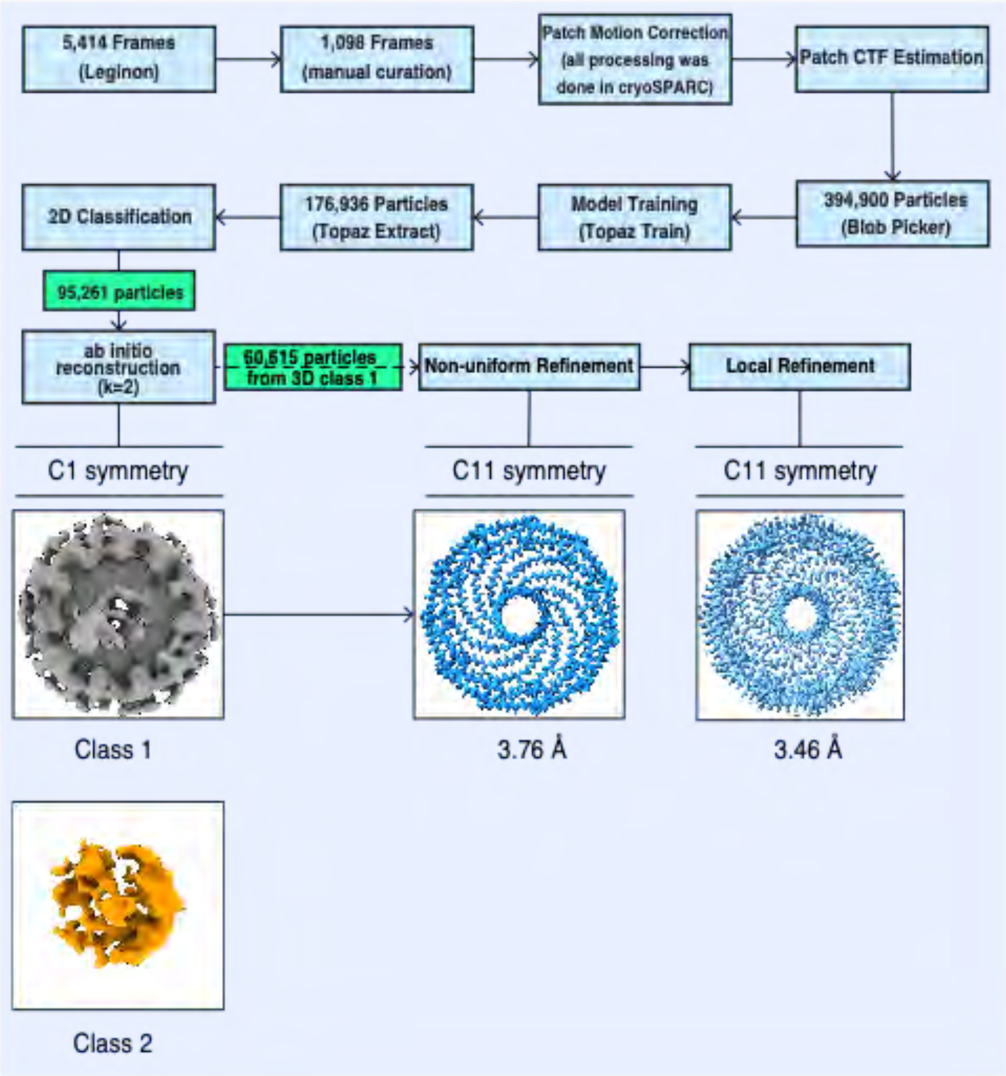

**Fig. S2. Flow chart of cryo-EM processing steps.** All data processing steps following data collection were carried out in cryoSPARC. Using Blob Picker with a subset of 100 micrographs, 47,892 particles were selected and used as input to generate a neural network model of the particles in the Topaz Train routine. Using the model of the particles as input, Topaz Extract was then run overall 984 micrographs, resulting in the identification of 176,936 particles. After the removal of the particles at the micrograph edges, 140,242 particles were extracted into 400 px x 400 px boxes and submitted to two successive rounds of 2D class averaging. After 2D classification, 95,261 particles remained that were then used as input for *ab initio* 3D reconstruction into two classes with no applied symmetry. One class containing 60,615 particles led to a 12 Å 3D structure that appeared to have some symmetrical features. This *ab initio* 3D model was then refined to 3.8 Å applying C11 symmetry using the non-uniform refinement routine. The refined particles were then submitted to a round of local refinement (with C11 symmetry enforced), producing the final 3D reconstruction at 3.5 Å resolution.

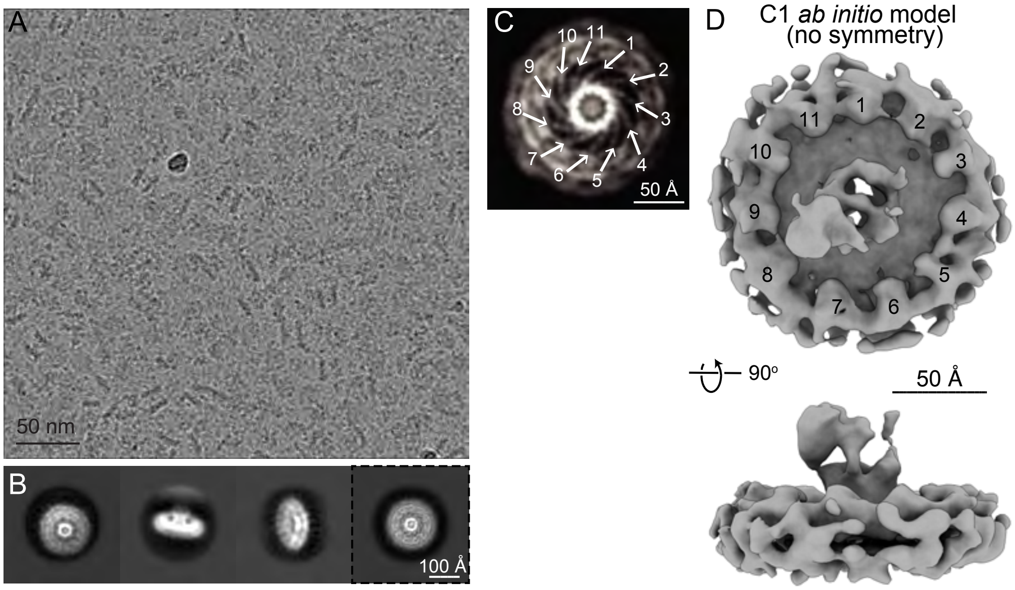

**Fig. S3. Single particle cryo-EM analysis of the 8S Cav1 complex. (A)** Motion corrected image of vitrified 8S Cav1 complexes. Scale bar, 50 nm. **(B)** Selected 2D class averages with no applied symmetry. Scale bar, 100 Å. 2D average with dashed line shown enlarged in panel C. **(C)** Enlarged 2D average with 11 $\alpha$-helices labeled. Scale bar, 50 Å. **(D)** *Ab initio* 3D reconstruction of the 8S Cav1 complex with no applied symmetry (C1). Structure rotated 90° around the x-axis. Position of the 11 subunits labeled in the *en face* view (top panel). Scale bar, 50 Å.

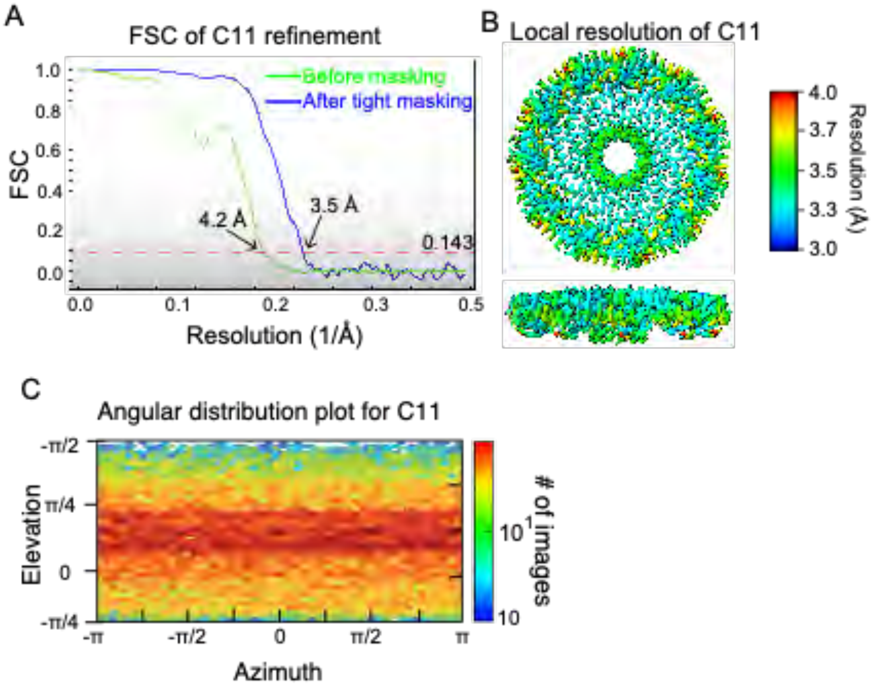

**Fig. S4. Cryo-EM processing of 8S Cav1 complex. (A)** Fourier Shell Correlation **(**FSC) of C11 refinement before (green line) and after (blue line) applying a tight mask in cryoSPARC. Dotted red line, Fourier Shell Correlation (FSC) =0.143. **(B)** Heat map showing local resolution of C11 3D reconstruction. **(C)** Euler angle plot of angles of particle distribution in the C11 3D reconstruction. The heat map scales are exponential.

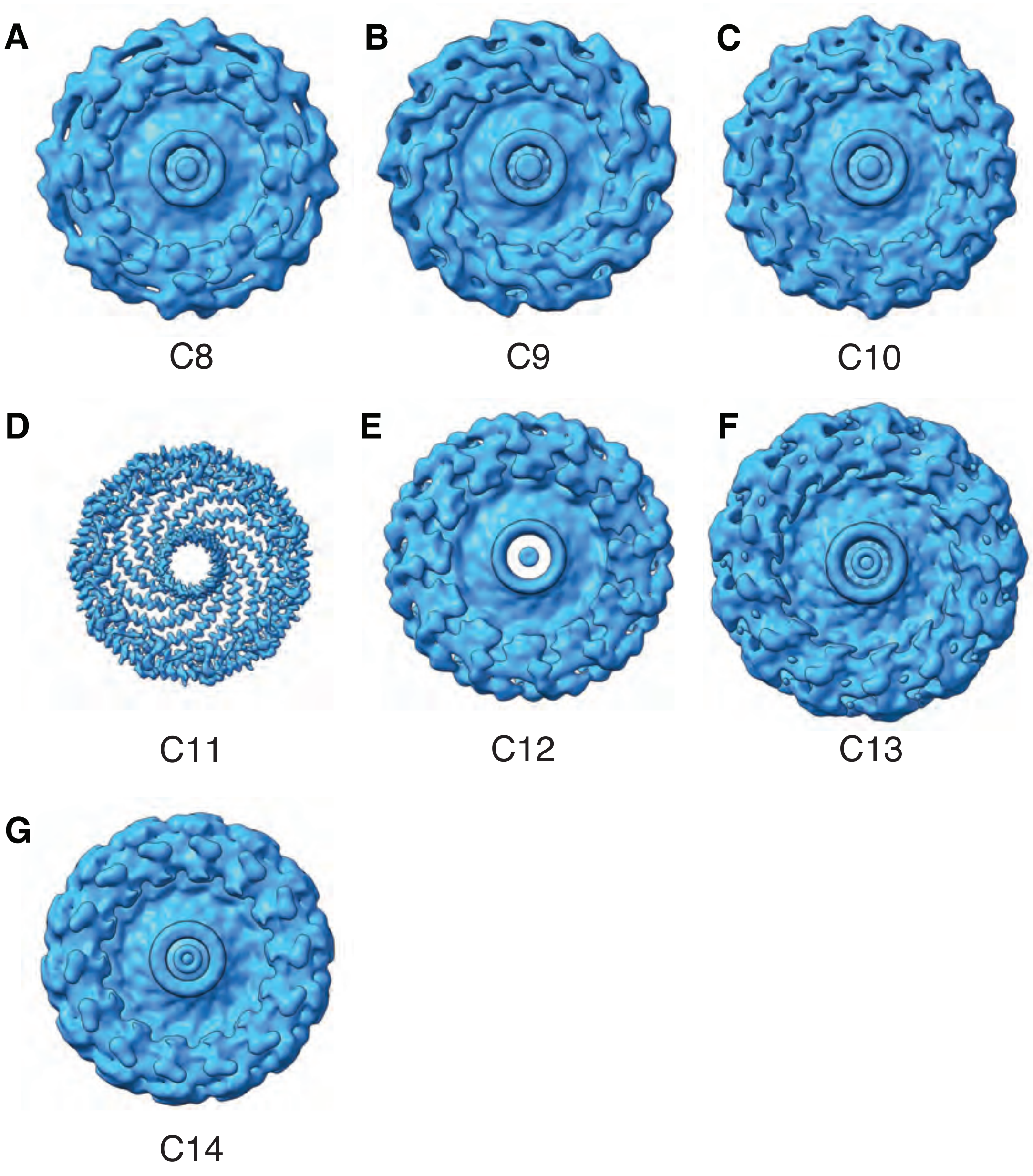

**Fig. S5. Applying different symmetries to the 3D refinement of the 8S Cav1 complex.** 3D reconstructions enforcing **(A)** C8, **(B)** C9, **(C)** C10, **(D)** C11, **(E)** C12, **(F)** C13, and **(G)** C14 symmetry operators. The map contour level was set to 0.4 in Chimera for each map.

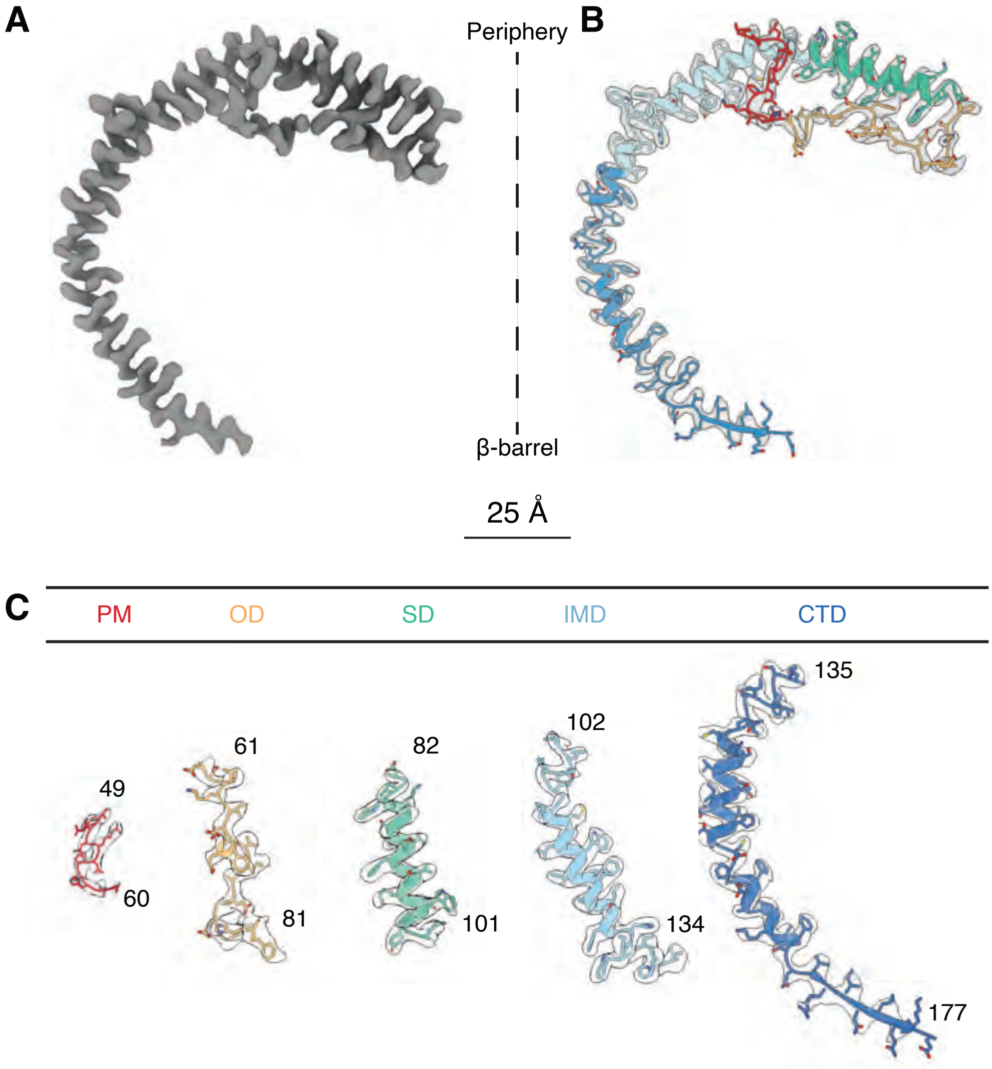

**Fig. S6. Examples of density fits of the Cav1 (49-177) atomic model. (A)** Density map for a Cav1 protomer (grey surface) extracted from the 8S complex structure. The protomer is oriented with the top being at the periphery of the 8S disc and the bottom the β-barrel. **(B)** Atomic coordinates (colored cartoon diagram) and fitted density map (transparent grey surface) for a single Cav1 protomer. Red, Pin Motif (PM); orange, Oligomerization Domain (OD); green, Scaffolding Domain (SD), light blue, Intermembrane Domain (IMD); dark blue, C-terminal Domain (CTD). **(C)** Coordinates of Cav1 regions fitted into the corresponding density maps (transparent grey surfaces) and their respective residue ranges. Domains are labeled above the structure. The contour level for the EM density map is at a value of 0.4 in Chimera.

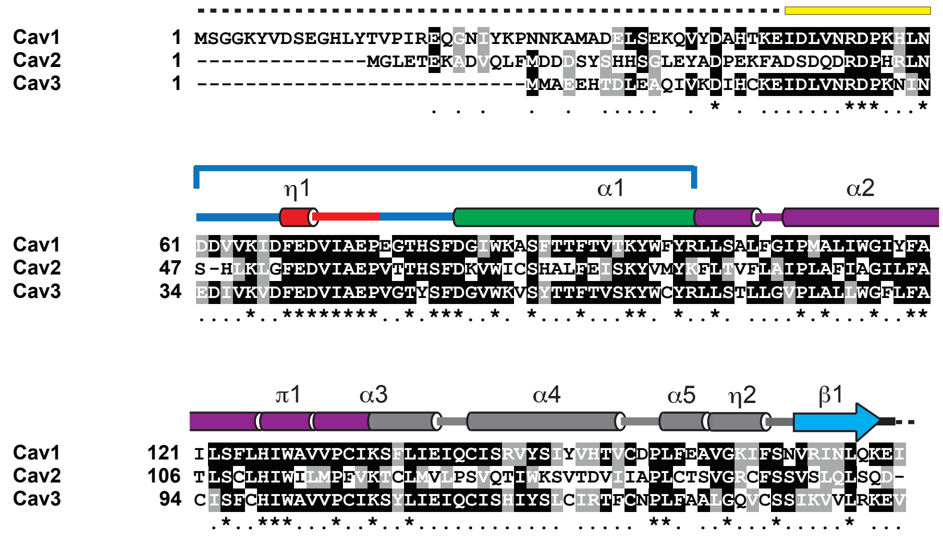

| Domains | Residues |
|---|---|
| PM: Pin motif | 49-60 |
| OD: Oligomerization domain | 61-101 |
| SM: Signature motif | 68-75 |
| SD: Scaffolding domain | 82-101 |
| IMD: Intramembrane domain | 102-134 |
| SR: Spoke-like region | 135-169 |
| β-strand | 170-176 |

**Fig. S7. Sequence alignment of caveolin family members, along with the secondary structural elements identified in the Cav1 structure**. Cylinders correspond to α-helices, arrow indicates β-strand, solid lines represent loops, and dashed lines are regions not resolved in the structure. The key domains are colored as in Figure 2, and color coding and residue range for individual domains are indicated at the bottom panel. The bracket shows the OD that encapsulates the SM and SD.

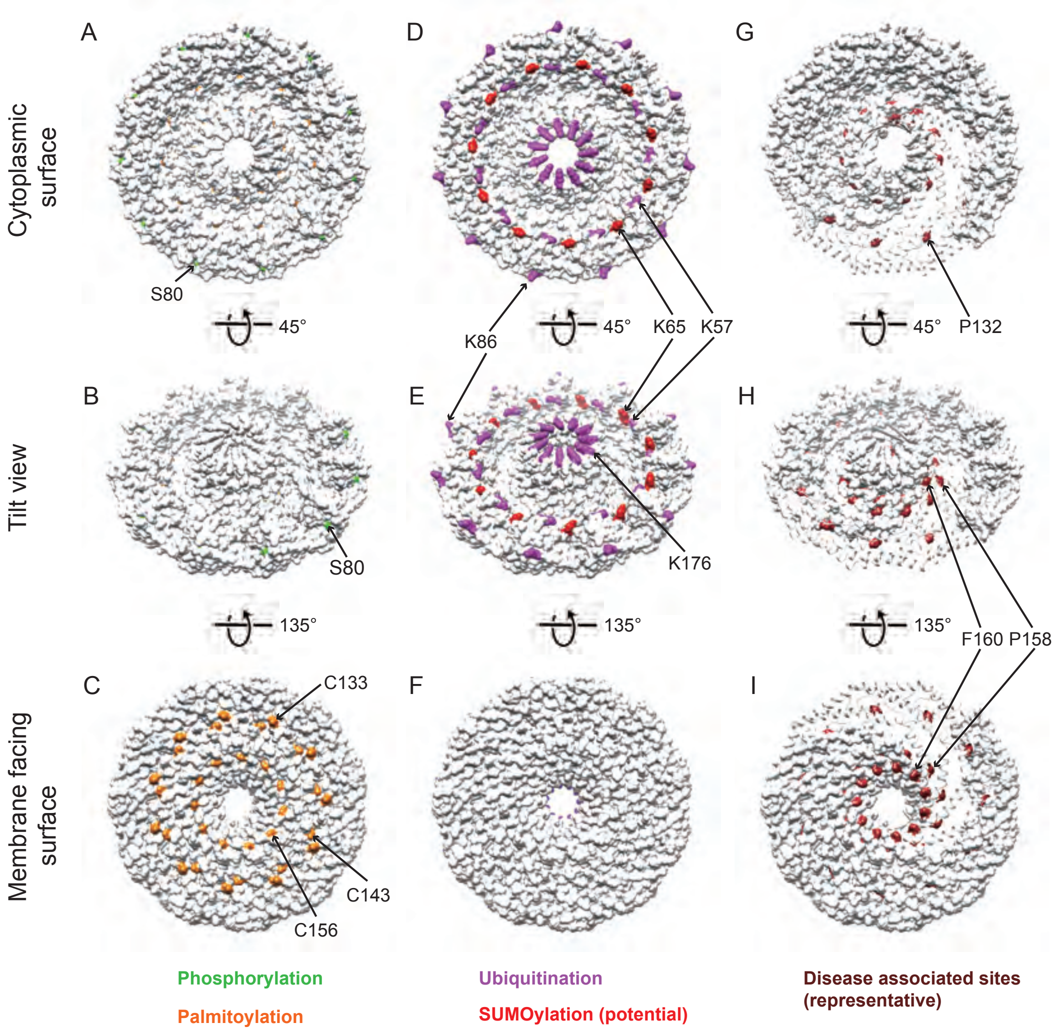

**Fig. S8. Location of Cav1 post-translational modifications and disease mutations on the 8S complex. (A-C)** Position of known phosphorylation and palmitoylation sites. **(D-F)** Position of known ubiquitination sites and potential SUMOylation sites. **(G-I)** Position of three disease associated mutations. Space filling models are shown in all panels except portions of panels G-I, where ribbon diagrams of several protomers were used to better illustrate the position of P132L.

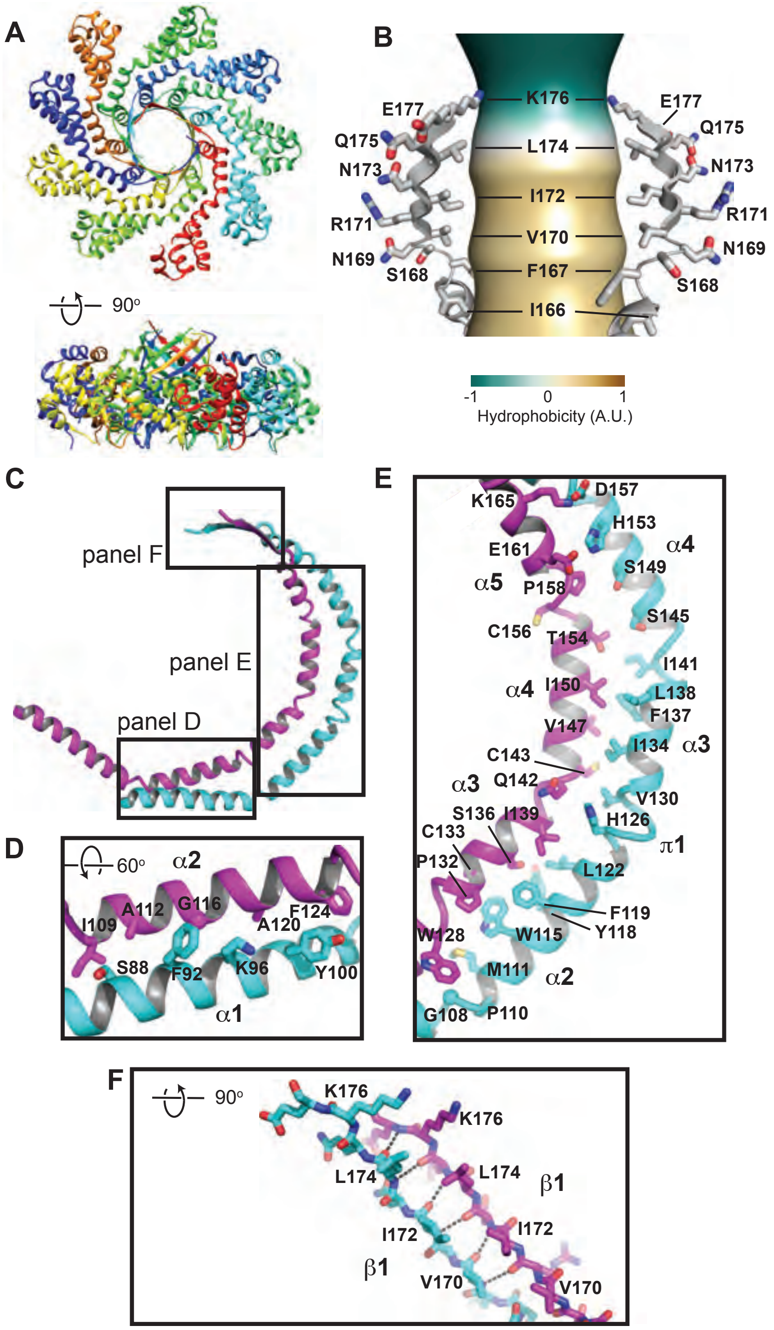

**Fig. S9. Features of Cav1's β-barrel and detailed views of the interactions between α-helices. (A)** The structure is from PDB ID 2AO9. (**B)** Close up view of the interior of Cav1's β-barrel. The pore size and hydrophobicity plot of the interior of the barrel were calculated using the software CHAP (*62*). Only two opposing protomers are shown for clarity. **(C- F)** Close up view of helix packing between two adjacent protomers highlighting residues lining the interfaces. Secondary structural elements are labeled. Protomers are depicted in two different colors (cyan and fuschia). Regions boxed in c are shown as zooms in d-f and are rotated by the indicated angles. **C.** Overview of two protomers, shown in an identical orientation as in Fig. 2C. N-terminal regions were omitted for clarity. **D.** Helical bundle crossing at the rim. **e.** Intramembrane domain and spoke region. **F.** Interactions between protomers in the spoke region extending to the β-strands of the barrel. Dashed lines indicate the hydrogen bonding between the backbone atoms of neighboring protomers at the β-barrel.

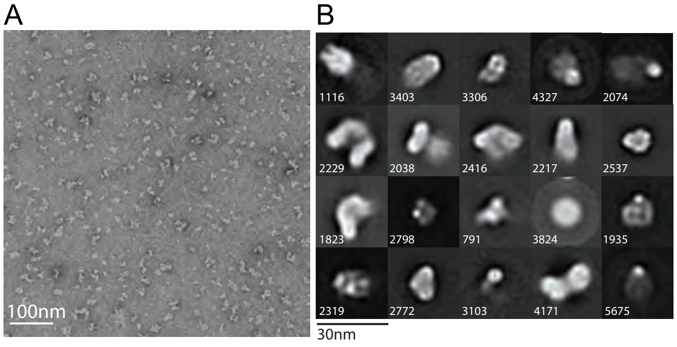

**Fig. S10. Cav1 R54A in the pin motif disrupts 8S complex stability. (A)** Representative negative stain image of Cav1 R54A. Scale bar, 100 nm. (**B)** 2D class averages. The number of particles in each class is shown in the bottom left corner. Scale bar, 30 nm.

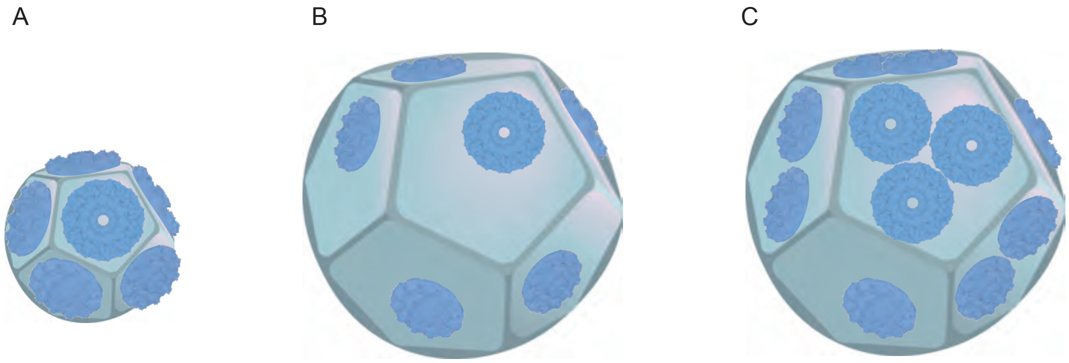

**Fig. S11. Packing of 8S complexes on dodecahedrons of the characteristic size of *h*-caveolae and caveolae.** Images are drawn to scale. **(A-C)** *h*-caveolae and caveolae are depicted as regular dodecahedrons assuming a circumradius of 15 nm for *h*-caveolae (*28*) and 30.5 nm for mammalian caveolae (*25*). 8S complexes are partially embedded in the membrane at a depth consistent with the model shown in Figure 4B. **A.** A single 8S complex fits on each face of a *h*-caveola. **B**. Representative packing density of 8S complexes on a mammalian caveola assuming a single 8S complex per face. **C**. Each face of a mammalian caveola can potentially fit up to 3 8S complexes.
